# *Azadirachta indica* (A. Juss.) reduces circulating *Dirofilaria immitis* microfilariae *in vivo*

**DOI:** 10.1101/781278

**Authors:** Niladri Mukherjee, Nikhilesh Joardar, Suprabhat Mukherjee, Santi P. Sinha Babu

**Affiliations:** Parasitology Laboratory, Department of Zoology (Centre for Advanced Studies), Visva-Bharati University, Santiniketan 731 235, West Bengal, India; Cancer Biology & Inflammatory Disorder Division, CSIR-Indian Institute of Chemical Biology, 4-Raja S.C. Mullick Road, Kolkata 700 032, West Bengal, India

**Keywords:** *Dirofilaria immitis*, *Azadirachta indica*, *in vivo*, Microfilaria reduction, *Canis familiaris*

## Abstract

The present study enumerates the effectiveness of ethanolic leaf extract of *A. indica* against circulating microfilariae (mf) of *D. immitis in vivo* of *Canis familiaris*. An ethanolic extract was prepared from the leaves of *A. indica* (EEA) and treated on dogs infected with the filarial nematode *Dirofilaria immitis*, the causative agent of canine cardiopulmonary dirofilariasis. Before treatment, all the infected dogs were vigilantly supervised for any natural fluctuation of mf count. Two doses; 25 and 50 mg/kg body weight/twice a day; both for 15 days were selected for *in vivo* tests along with a control group which received an empty capsule during the study period. The highest reduction of circulating mf was counted on day 60, showing mf reduction of 77.9% and 86.7% respectively for the two doses. Thereafter mf density increased with a minor change and maintained reduction of 49.5% and 64.1% on day 180 respectively. Additionally, no appreciable side effects in the treated dogs were recorded as evident from serum toxicity parameter analyses. In conclusion, the ethanolic formulation of *A. indica* leaves possesses considerable effectiveness against *D. immitis in vivo* with no toxic modification in the host after exposure. Thus, EEA appears to be a good alternative remedy against heartworm infection in infected dogs.

## 1. Introduction

Cardiopulmonary dirofilariasis or heartworm disease of dogs and other canids is caused by the filarial nematode *Dirofilaria immitis* (*D. immitis*, family Onchocercidae) and widely distributed in the subcontinents of Africa, Australia, Asia and Europe (Genchi et al., 2005). The causative parasite naturally resides in the pulmonary arteries, occasionally inside the right heart and sometimes the aberrant movement of them to various other parts of the body of affected animals can be noted (Bowman et al., 2009). Diversified clinical symptoms due to *D. immitis* infection in dogs associated with different stages of infection that can ultimately lead to the death of the affected animals (Bowman et al., 2009, Nelson et al., 2014). Adult females upon copulation release thousands of microfilariae (mf) into the circulation and make it prone for the transmission through mosquitoes. Infection to naive canids occurs when infective (3^rd^ stage that developed from mf after two molting inside the vector) larvae of *D. immitis* enter into the circulation via a vector (Bowman et al., 2009). Treatment against heartworm disease is troublesome, as melarsomine dihydrochloride, the only approved macro-filaricidal by the US FDA having some complications in infected dogs as the dead worms decompose into small fragments which can cause blockage of the pulmonary arterioles and capillary beds that can lower the blood flow significantly and carries risk of probable lethality with some adverse neurological reactions (Godel et al., 2012). Macrocyclic lactones (ivermectin, milbemycin oxime, selamectin, moxidectin) are in use for more than 25 years as microfilaricides that mainly block transmission (Godel et al., 2012, Bourguinat et al., 2011). However, resistance of *D. immitis* against these choices of drugs is definitely a possibility and a matter of discussion and it was found that almost 1/5^th^ of the dogs that received periodic heartworm preventive medicine of the macrocyclic lactone groups without any adulticidal agents can be continued to have circulating microfilariae that possibly can evolve into resistance strain (Bowman and Mannella, 2011). Thus, alternative therapeutics is essentially needed. In this regard, some plant extracts or isolated phytocompounds have shown potentials to become new alternative chemotherapeutics against heartworm disease. In our previous research it was found that ethanolic extract from the funicles of *Acacia auriculiformis* and from the leaves of *Azadirachta indica* was found to be effective against *D. immitis, in vivo* and *in vitro*, respectively (Chakraborty et al., 1995, Mukherjee et al., 2014). Previously, we have demonstrated the antifilarial activity of an ethanolic extract from the leaves of *A. indica* (EEA) on *D. immitis in vitro* through alteration of redox parameters and subsequent apoptosis induction (Mukherjee et al., 2014). Here, we extended our previous study by examining the effectiveness of EEA against circulating *D. immitis* mf *in vivo* using the natural host of the parasite i.e. *Canis familiaris*.

## 2. Materials and Methods

The protocol of this study was approved by the Institutional Animal Ethical Committee, Visva-Bharati University, Santiniketan– 731 235, India. Animal sampling was performed under the surveillance and guidance of trained officials and resource persons from Bolpur Sub-divisional Veterinary Hospital, Department of Animal Resource Development, Govt. of West Bengal, India. Adult dogs of both male and female were included in the study apart from the injured, lactating or pregnant ones. Ethanolic extract of the leaves of *A. indica* was prepared and characterized for chemo-profiling according to Mukherjee et al (2014). Various doses (25 and 50 mg/kg b. wt. / twice a day) of EEA were selected for treating the infected dogs orally and were prepared in empty capsules. Before treatment, *D. immitis* infected adult street-dogs (*C. familiaris*) were screened by random sampling of blood withdrawn from the recurrent tarsal vein of the hind legs with the aid of sterile Vacutainer equipment (4ml, plastic whole blood tube with Spray-coated K2EDTA; BD, USA) in their natural territorial zones (all the sampling were done at 9 A.M. ± 15 min IST). Circulating mf were detected by visualizing the Giemsa stained blood slide under the microscope following our previous report (Mukherjee et al., 2014). For *in vivo* studies, the infected dogs were randomized into control and experimental groups and distributed into 3 groups of animals (n= 6) as shown in Table 1. For each group prior to *in vivo* trial, continuous blood sampling was performed at weekly intervals for 12 weeks to observe any natural change over time (Table 2). Group II and Group III, received EEA at 25 and 50 mg/kg b.wt./twice a day, respectively. Group I, served as control (matching placebo for 15 days). EEA was given for 15 days continuously in empty capsules filled with it and embedded in a piece of bread at 12 h interval. To observe the changes in the circulating *D. immitis* mf count due to EEA administration continuous sampling was carried out for 30 days with a weekly interval thereafter at 15 days interval up to day 90. Additional sampling was performed thereafter with a monthly interval of up to 6 months. Evaluation of toxicity due to EEA exposure was performed following the guidelines from the Organization for Economic Cooperation and Development (OECD) (OECD, 2009). Biochemical tests included measurement of serum total protein, albumin, globulin, total bilirubin both conjugated and un-conjugated, serum glutamate oxaloacetic transaminase (SGOT), serum glutamate pyruvic transaminase (SGPT) and alkaline phosphatase following the standard laboratory methods at different time points during treatment and posttreatment, i.e., day 0, 15, 30 and 180. Data were collected in triplicates, subjected to Student’s t-test after accessing normality distribution through Shapiro-Wilk test of normality by SPSS software and finally expressed as mean ± SEM. Significant differences, p-value <0.05 (*), <0.001(**) between the means of each sample group on the day ‘0’ count and experiment day count were analyzed by using MS Excel.

**Table 1.**
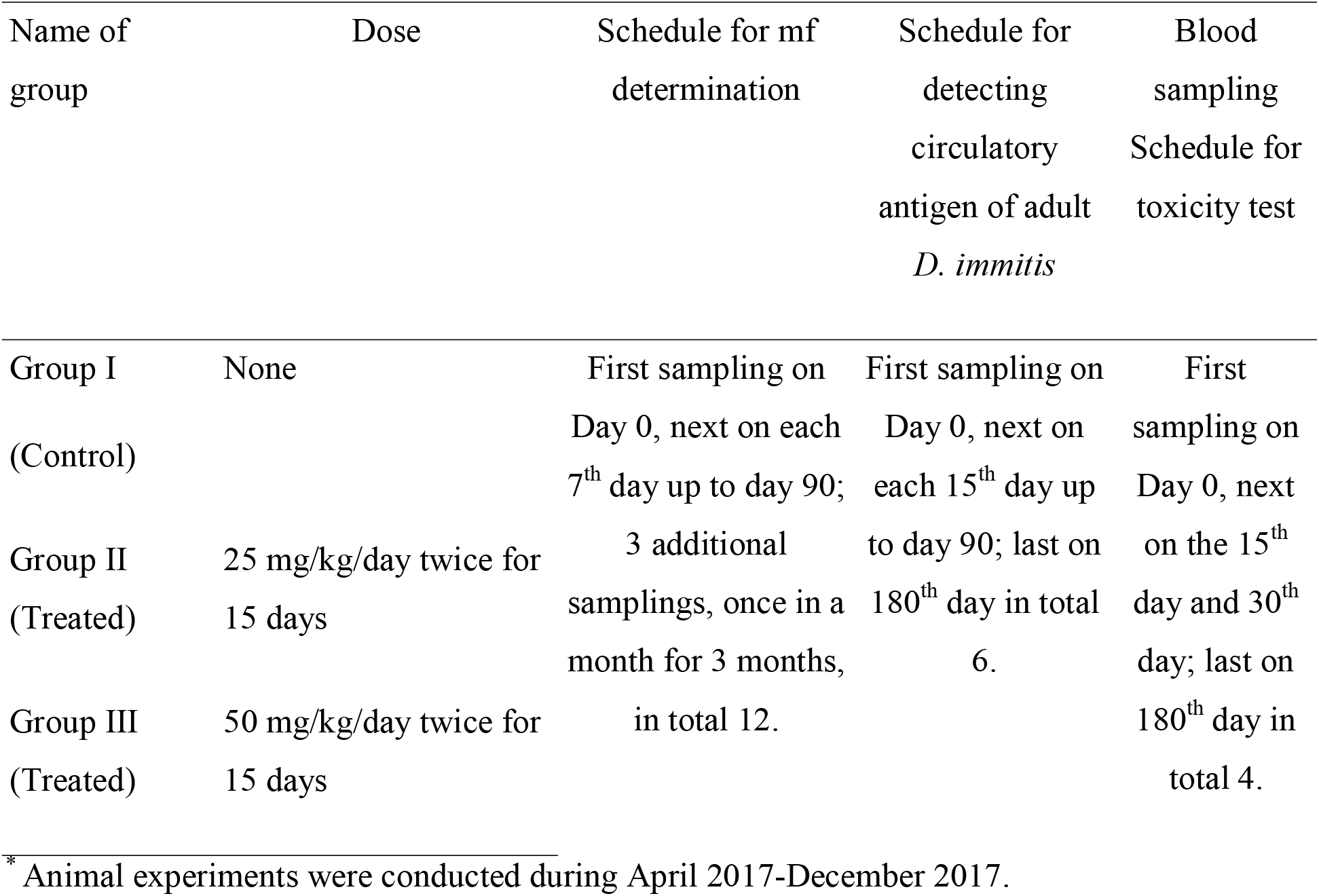
Schedule* for *in vivo* experiments and assessment of side effect

## 3. Results and Discussion

The traditional rather ethno-medication system still has the potential to serve in the modern medicinal practice especially at drug developmental research as apart from their direct effectiveness against different ailments, natural products can even provide a storehouse of various active compounds. Primary evaluation of the efficacy of EEA against *D. immitis in vitro* showed that EEA caused its effectiveness by altering the redox parameters and subsequent induction of apoptosis in *D. immitis* mf (Mukherjee et al., 2014) and therefore EEA have possibilities of controlling mf densities in infected animals and act as a transmission regulator. Random microscopic examinations for *D. immitis* mf infections of dogs revealed that infections were at different densities and thus grouped into three with six animals in each group having comparable infections and did not vary significantly during pretreatment period (Table 2; Fig. 1-A). Treatment results were provided in Table 2 and Fig. 1-B and Fig. 1-C. Day 0 count is the mean of all the pretreatment observations. Oral administrations of EEA at 25 and 50 mg/kg b.wt./twice a day for 15 days revealed that EEA was effective in reducing circulating mf count significantly and that too was dose-dependent. For Group II (25 mg/kg b.wt./twice a day for 15 days) EEA treatment resulted in a reduction in mf density and the maximum reduction of mf count (77.9%) was recorded on day 60 relative to pre-treatment count. Thereafter mf density increased with a minor change and maintained reduction of 49.5% on day 180. Whereas for group III (50 mg/kg b.wt./twice a day for 15 days), mf count was reduced drastically to 86.7% on day 60 but like that of the group II got increased thereafter and finally maintained 64.1% reduction on day 180. Significant (p< 0.05) reduction of circulating mf as well as percentage reduction were clearly dose-dependent and followed a particular trend, reached maximum reduction on day 60 and then got increased but the increment of mf count post-treatment was much slower than initial reduction in circulating mf count after EEA administration (Fig. 1-B and Fig. 1-C). The effect of EEA on circulating mf did not show a drastic fall in mf count suddenly, but the effect was continuous and persistent. The continuous depletion is more useful as a sudden mass death of mf can cause severe lethal complications due to the sudden heavy release of mf antigens (Merawin et al., 2010). To check if EEA has any effect on adults, from each blood sample, serum was also collected and examined for adult *D. immitis* Ag (Antigen Rapid Canine Heartworm Ag Test Kit 2.0” BioNote, Inc. Korea). It was found that EEA did not produce any considerable change in circulating adult Ag repertoire and possibly EEA produced little or no effect on adult parasites (Fig. 1-D). Preventive action of EEA executed through altruistic prevention as reduction of the circulatory mf was highly evident but the persistence of the adult worms in the dogs. It was noticed that EEA may not have any profound effect on the adults of *D. immitis*, as noticed by the antigen detection tests but reduction in circulating mf can have an effect on transmission blockage which is in continuation of our earlier report where EEA mediated apoptotic death of the exposed *D. immitis* mf in vitro (Mukherjee et al., 2014). Although it is very early to emphasize this part of the conclusion as the remaining number of mf is quite enough to continue the transmission but opens provision of further research and finding out the most important compound from EEA that is actually causing the reduction and maybe in future can confirm a therapeutic lead as an complete circulatory *D. immitis* macrofilaricidal agent. Furthermore, EEA administration did not alter any of the physico-biochemical parameters significantly alongside the improvement of liver function in the treated dogs by means of sustained reduction of serum bilirubin level, alkaline phosphatase, SGOT and SGPT were also noticed (Table 3).

**Fig. 1.**
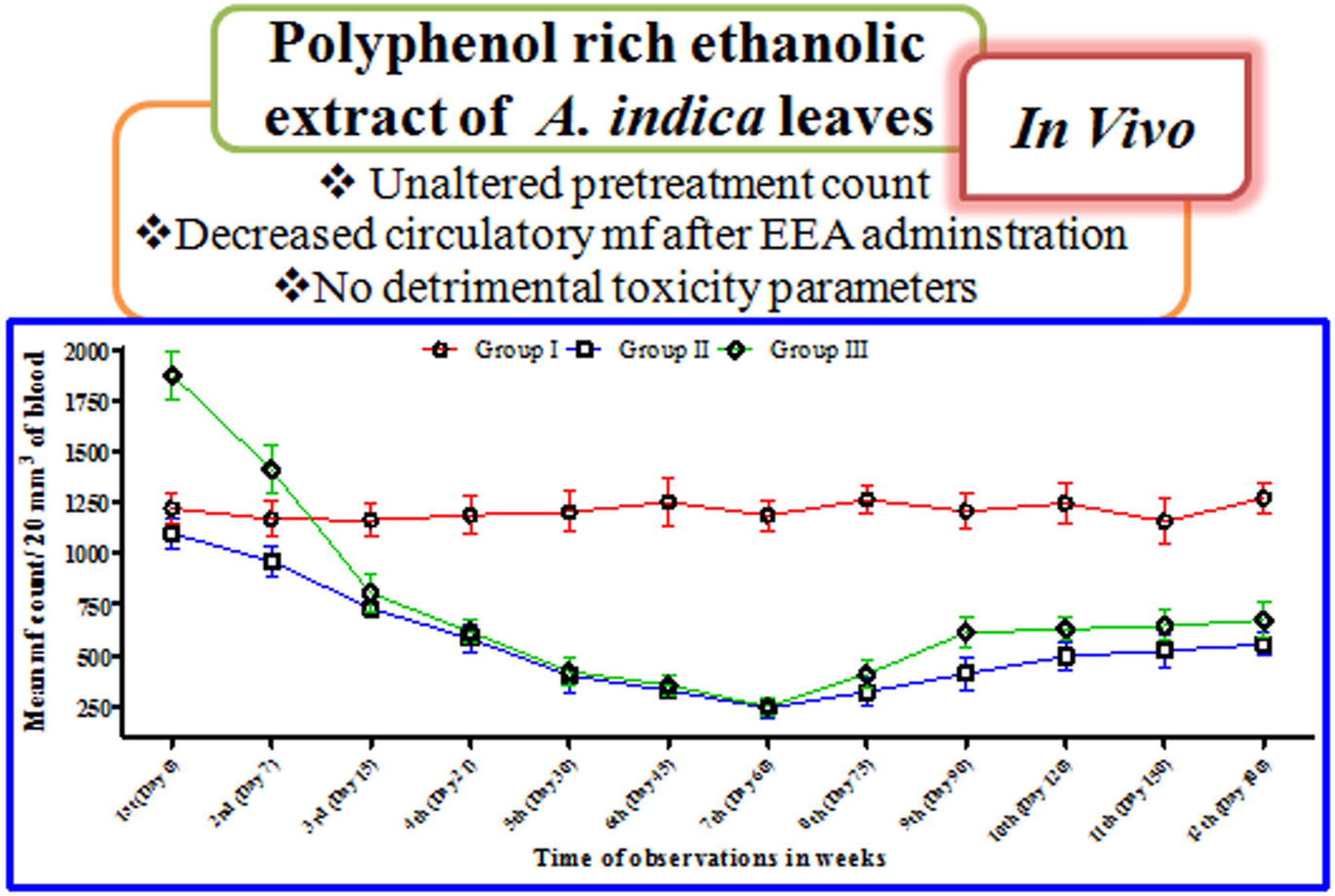
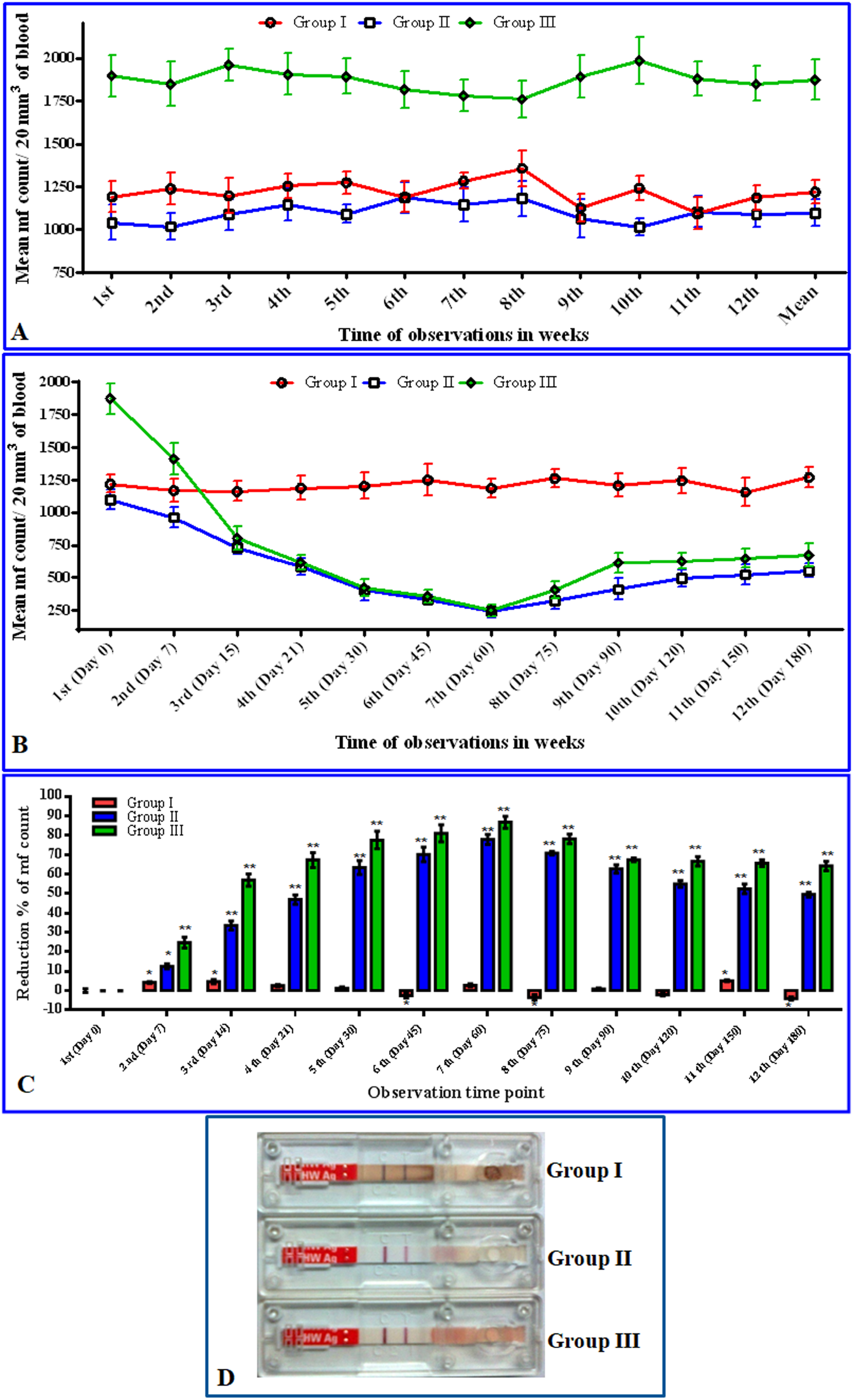
*In vivo* activity of EEA. (A) The *D. immitis* mf load in the experimental dog groups did not vary appreciably during the pretreatment period as counted for twelve consecutive weeks. (B) The changes in mf count at a different time of observations are plotted in case of the treated and control (matching placebo) groups. In case of treated groups (25 and 50 mg/kg b.wt./twice a day for 15 days) circulating mf count declined significantly in a dose-dependent manner. A curve was obtained which showed the declining pattern of microfilarial count at different time intervals. Continuous reductions of mf count were noticed up to day 60 but thereafter mf density had minor up-regulation. (C) The percent reduction in mf count following treatment with EEA in comparison to control at various time intervals. For both, the groups’ highest reductions were noticed at day 60 and were about 77.9% and 86.7% for group II and III respectively. (D) EEA was unable to produce any considerable change in circulating adult Ag repertoire after treatment schedule for control Group I and treated groups, Group II and III. Each bar represents the mean ± SEM. Data were analyzed by paired‘t’ test using MS Excel software. There was a significant difference between the control and the treated groups [p< 0.05 (*) and < 0.001(**) considered as statistically significant].

**Table 2.**
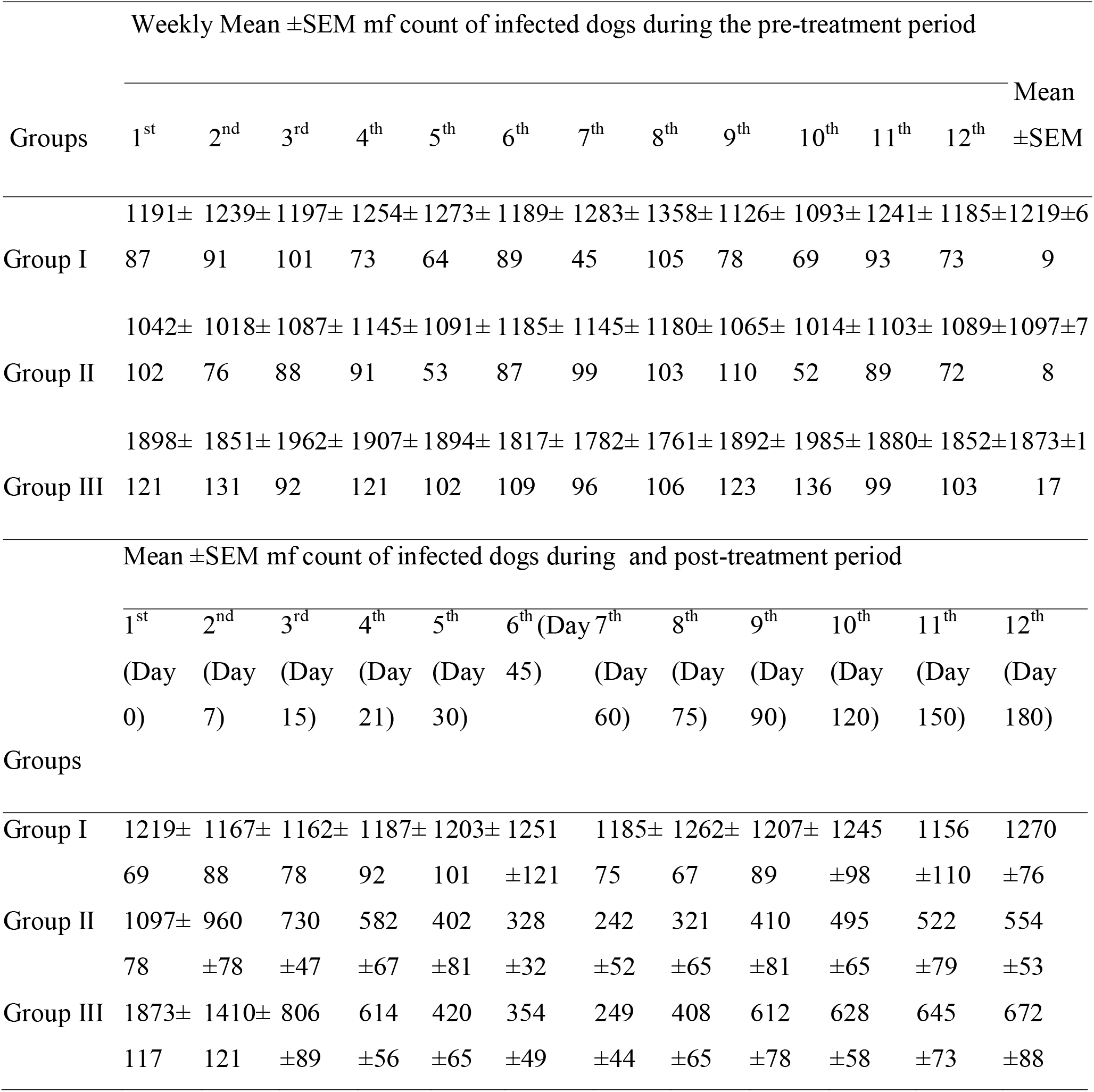
Microfilariae (mf) count per 20 mm^3^ blood of dogs naturally infected with *D. immitis* prior to treatment; during and after treatment

**Table 3.**
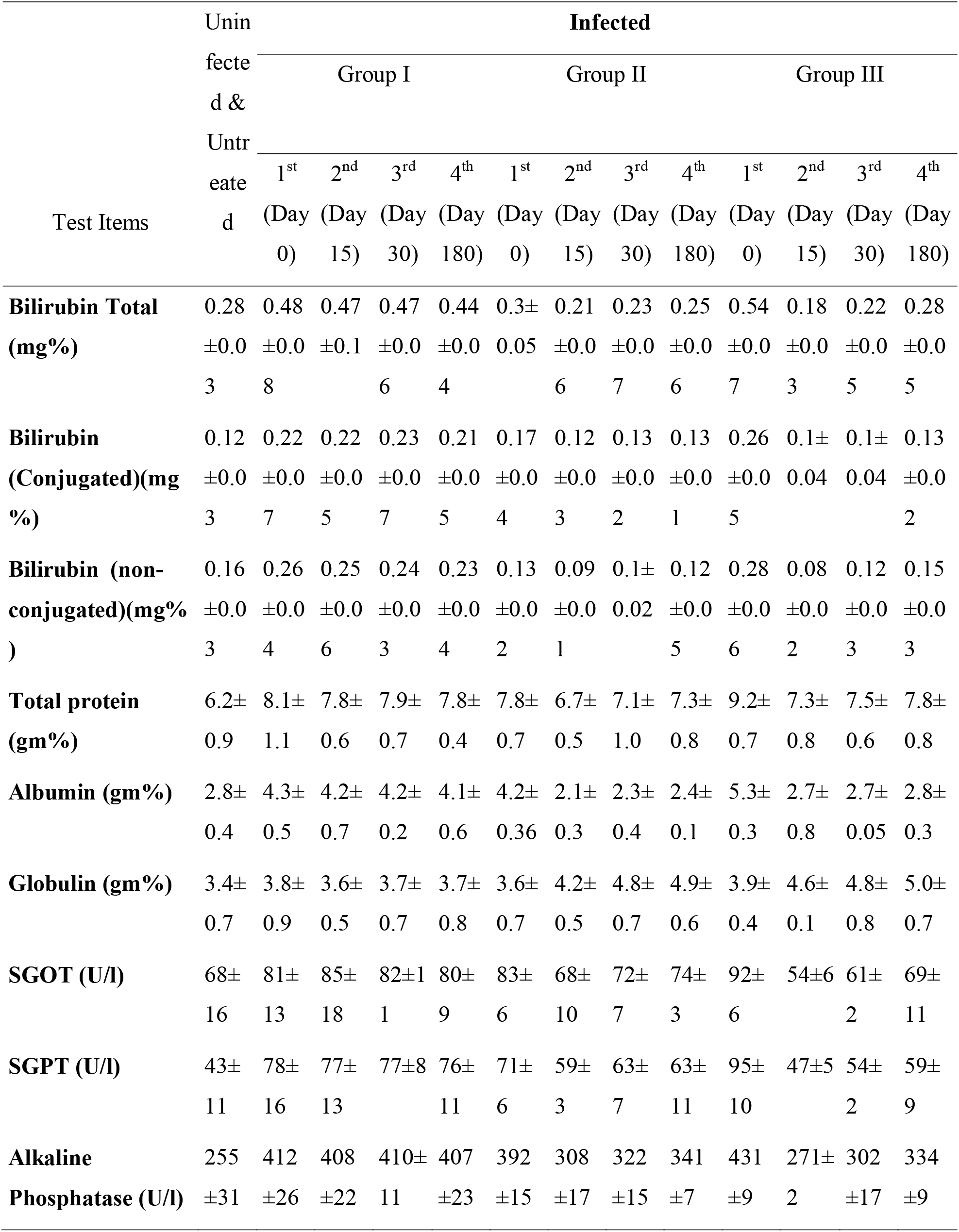
Toxicity profiles of EEA in dogs as revealed by serological parameters

## 4. Conclusion

Transmission of *D. immitis* mf does majorly reliant on their sufficient presence in the host circulation as well as on the number of infected hosts where vector availability is not of any question. Therefore, the use of EEA suppresses mf count in the infected population and additionally improves the overall health status of treated dogs which can be helpful to combat the disease. With additional research and exploration of the main effective compound of the extract liable for circulating microfilaria reduction can emerge as an alternative mean during unavailability of the common microfilaricides and during the occurrence of resistance. Considering our overall findings, EEA appears to evolve as a possible remedy to control *D. immitis* in a safe, cost-effective and efficacious way.

## Acknowledgments

We thank the University Grants Commission (UGC) (Grant No. 42-534/2013 (SR) Govt. of India for supporting this work financially. The manuscript has been checked critically by the full version of the Grammarly software.

## Ethical statement

The protocol of this study was approved by the Institutional Animal Ethical Committee, Visva-Bharati University, Santiniketan-731 235, India. Animal sampling was performed under the surveillance and guidance of trained officials and resource persons from Bolpur Sub-divisional Veterinary Hospital, Department of Animal Resource Development, Govt. of West Bengal, India.

